# CaMKII T286 autophosphorylation is not propagated at basal Ca^2+^ levels and is required only for the induction phase of LTP

**DOI:** 10.64898/2026.07.27.741023

**Authors:** Fan-Yi Chao, Nicole L. Rumian, Adam P. Miller, Steven J. Coultrap, Steve L. Reichow, K. Ulrich Bayer

**Author notes:** Correspondence should be addressed to Steve L. Reichow or K. Ulrich Bayer.

## Abstract

The Ca^2+^/calmodulin-dependent protein kinase II (CaMKII) and its autophosphorylation at T286 (P-T286) mediates long-term potentiation (LTP) of synaptic strength. CaMKII forms stable 12-meric holoenzymes, but exchange of its subunits is proposed to propagate the P-T286 state beyond dephosphorylation or degradation of individual subunits. However, a recent study questioned subunit exchange and instead proposed P-T286 propagation by *trans*-holoenzyme phosphorylation. We show here that both subunit exchange and *trans*-holoenzyme P-T286 can occur (although *cis*-holoenzyme P-T286 is much preferred), but that neither mechanism propagates P-T286 at basal cellular Ca^2+^ levels (50-100 nM) in absence of further Ca^2+^-stimuli. Functionally, we show that after theta-burst stimulation (TBS), P-T286 is required only during the first minutes of LTP induction that prime for subsequent LTP expression, but neither for LTP expression itself nor for LTP maintenance. These results indicate that P-T286 of CaMKII does not self-perpetuate and does not mediate storage but computation of synaptic information.

## INTRODUCTION

Learning and memory are thought to require forms of synaptic plasticity, specifically including hippocampal long-term potentiation and depression (LTP and LTD), which both require the brain-specific Ca^2+^/calmodulin-dependent protein kinase II α isoform (CaMKIIα; here referred to as CaMKII for short) and its autophosphorylation at T286 (P-T286)^1,2^ (as recently reviewed^3–5^). This P-T286 is thought to occur between two subunits within the 12-meric CaMKII holoenzyme^6,7^ and has long been regarded as a form of “molecular memory”, as it generates Ca^2+^-independent “autonomous” activity of CaMKII that outlast the initial Ca^2+^-stimulus^8,9^, thereby enabling CaMKII to “remember” past Ca^2+^-stimuli. These properties gave rise to the hypothesis that CaMKII stimulation by Ca^2+^/calmodulin is part of the information processing that leads to LTP induction, whereas P-T286-induced autonomous CaMKII activity provides the information storage that enables LTP maintenance. This hypothesis has remained highly attractive (as previously reviewed^5,10^) despite strong arguments and experimental evidence to the contrary^11–14^ (as also reviewed previously^5,15^). For instance, a recent electrophysiological study claimed that its finding of maintained synaptic strength requires maintenance of P-T286 beyond protein turnover^16^ by one of two possible mechanisms: subunit exchange within the CaMKII holoenzyme^17,18^ or a *trans*-holoenzyme mechanism of the P-T286 reaction^19^. Thus, we decided to re-examine both of these two possible mechanisms, especially since the subunit exchange between holoenzymes has recently been questioned^19^. Additionally, we re-examined the requirement of sustained P-T286 for LTP maintenance after induction by theta-burst stimulation (TBS).

First, we corroborated the original experiments that had provided the initial suggestion for a possible subunit exchange^20^. Then, we corroborated that P-T826 can indeed occur by a *trans-*holoenzyme P-T286 reaction (that occurs between subunits of two different holoenzymes), although the *cis*-holoenzyme reaction (that occurs between the subunits within one holoenzyme) was much more favored. However, we also show that neither of these two mechanisms can propagate P-T286 in absence of Ca^2+^ or at the low levels of basal cellular Ca^2+^ levels (50-100 nM), as both the *trans*- and the *cis*-holoenzyme P-T286 required higher levels of Ca^2+^ stimulation. These findings are consistent with a model in which P-T286 mediates synaptic information processing, not storage^5,15^. Indeed, functional experiments in slices from the P-T286-incompetent CaMKII T286A mutant mice further supports that the storage of synaptic information that is mediated by LTP maintenance does not require P-T286 at all.

## RESULTS

### Probing for subunit exchange within the CaMKII holoenzyme after partial tryptic digest

The initial indication for subunit exchange came through partial proteolytic cleavage experiments^20^. Direct structural comparisons by electron microscopy (EM) indicated that full-length CaMKII holoenzymes exist primarily as dodecameric (12-mer) complexes (Figure 1A), whereas isolated association domains (blue) lacking kinase domains (yellow) preferentially adopted tetradecameric (14-mer) assemblies^20^. Notably, when the 12-meric full-length holoenzymes were subjected to a partial tryptic digest that removed the kinase domains, analysis of the remaining association-domain assemblies indicated transformation to 14-mers, suggesting that subunit exchange and hub reorganization must have occurred. However, while the pre-digest assemblies were analyzed by EM, these 14-meric post-digest assemblies were instead detected by X-ray crystallography^20^, leaving the possibility for preferential crystallization of 14-mer over 12-mer association-domain assemblies^21^ and potential overestimation of the 14-mer fraction. Specifically, if the sample contained a mixture of 12-mer and 14-mer holoenzymes already prior to proteolysis, it is possible that the 14-mer association-domain assemblies generated by proteolysis from the 14-mer holoenzymes could preferentially crystallize, even if the 12-mer assemblies remained the predominant population in solution. Consistent with this possibility, our previous EM analysis of full-length CaMKIIα holoenzymes identified predominantly 12-mer assemblies but also detected a minor population (<4%) of 14-mers containing kinase domains^22^. Notably, other preparations have been reported to contain approximately equal populations of 12-mer and 14-mer assemblies^18^, although it remains unclear whether kinase domains remained associated with the majority of those 14-mer particles. To eliminate the potential bias introduced by crystallization methods, we repeated the partial tryptic digest experiment using our previously characterized CaMKIIα preparation protocols and directly assessed the resulting oligomeric distributions by negative-stain EM.

**Figure 1.**
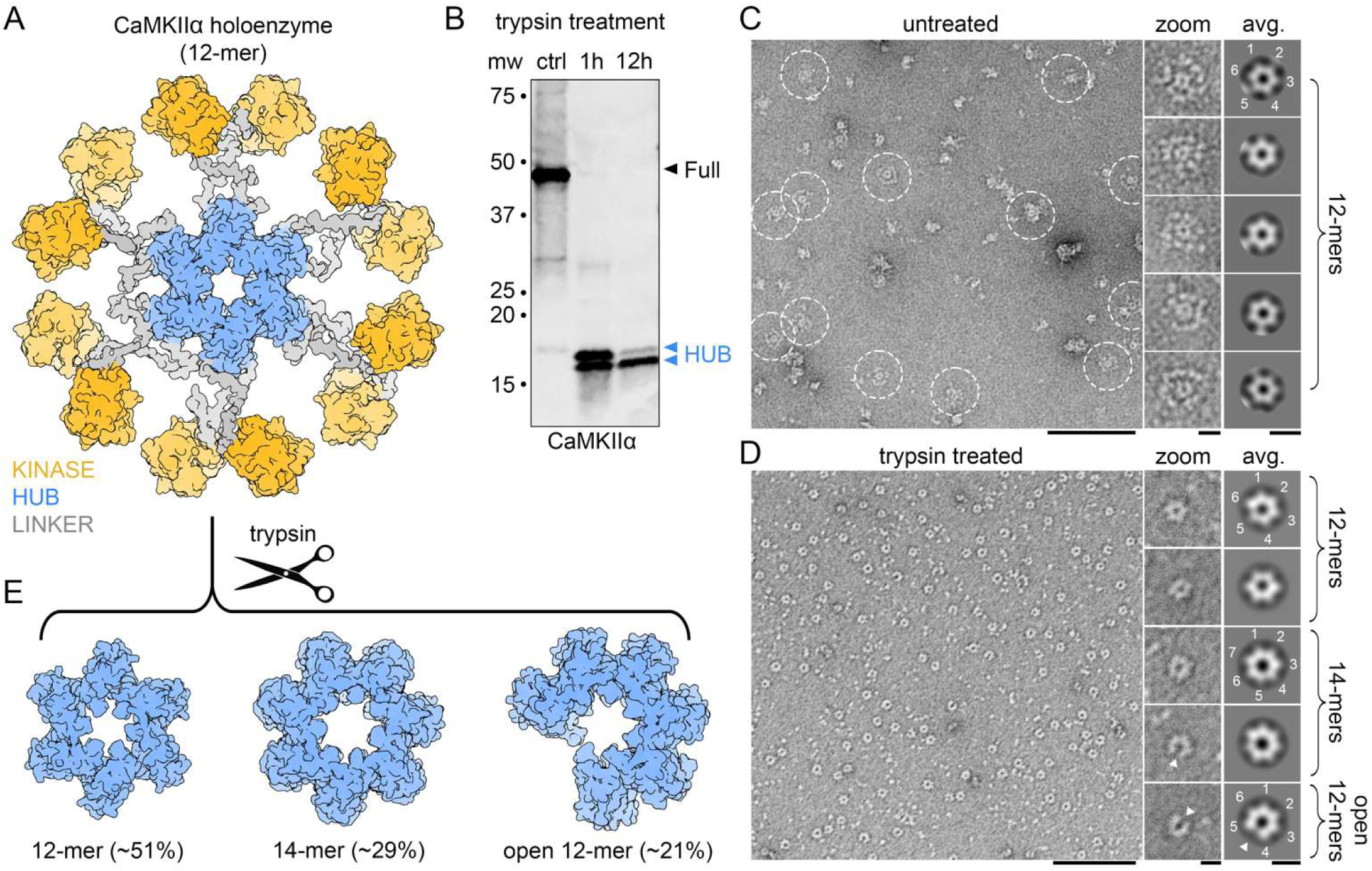
Partial tryptic digestion of CaMKIIα holoenzymes promotes hub remodeling and formation of alternative oligomeric states. (A) Structural model of the 12-meric CaMKIIα holoenzyme (PDB 5U6Y) showing the central hub domain (blue), kinase domains (KD; yellow), and connecting linker regions (gray). (B) SDS–PAGE and western blot analysis demonstrating limited proteolysis by trypsin removes the kinase and linker regions while leaving the association-domain hub intact. The digest was stopped with PMSF either after 1 h (with 11 h additional incubation) or after 12 h. (C,D) Negative-stain EM analysis of untreated CaMKIIα holoenzymes *(C)* and trypsin-treated samples *(D)*. *Left*, representative micrographs. *Middle*, zoom-views of representative particles. *Right*, representative 2D class averages following focused classification of the hub domain. Untreated particles display canonical holoenzymes with a central hub surrounded by peripheral kinase domain densities and classify exclusively as 6-fold symmetric 12-mers. In contrast, trypsin-treated particles appear predominantly as isolated hub assemblies and classify into three major populations corresponding to 12-mers, 14-mers (7-fold symmetry), and partially opened 12-mers (“open 12-mers”). Triangles in *(D)* indicate the site of opening of the hub assembly. Scale bars shown for micrographs = 100 nm and for inset and class averages = 10 nm. (E) Structural models corresponding to the three hub-domain architectures observed following trypsin treatment: 12-mer (PDB 5IG3), 14-mer (PDB 1HKX), and open 12-mer (PDB 8T6Q). Approximate populations for each assembly resolved by EM are indicated.

Successful partial proteolysis liberating the kinase domains while leaving the association domains intact after trypsin treatment was verified by western blot analysis (Figure 1B and S1). Without trypsin treatment, CaMKIIα holoenzymes displayed canonical sixfold symmetric hub assemblies surrounded by peripheral kinase domains, consistent with our previous EM studies^22,23^ (Figure 1C). Focused classification of hub regions resolved exclusively 12-mer assemblies in the untreated dataset (n = 1,396 particles; Figure 1C, right). The absence of detectable 14-mer classes in the current control dataset likely reflects the smaller particle number analyzed here (1,396 versus 8,081 particles previously reported), limiting classification of low-occupancy states.

In contrast, trypsinized particles appeared predominantly as isolated hub assemblies lacking peripheral kinase domain densities in raw micrographs (Figure 1D). Following 1 h trypsinization, the reaction was quenched with PMSF and further incubated overnight at 25°C to allow equilibration of subunit exchange dynamics. Focused classification of these proteolyzed particles revealed a heterogeneous distribution consisting of 12-mer (50.6%), 14-mer (28.6%), and partially opened sixfold symmetric “open 12-mer” hub assemblies that may represent a transition state (20.8%) (n = 1,515 particles; Figure 1D,E). Prolonged overnight exposure to trypsin produced qualitatively similar results (Figure S2). These observations indicate that proteolytic disruption of the holoenzyme promotes hub remodeling and redistribution by subunit exchange into alternative oligomeric states, while maintaining dodecameric assemblies as the predominant species.

Together, these observations now directly demonstrate by EM that trypsin-mediated proteolysis redistributes CaMKIIα hub assemblies from a predominantly monodisperse 12-mer population toward a more heterogeneous ensemble containing a significant fraction of 14-mers, thus supporting the notion of subunit exchange between CaMKII holoenzyme assemblies.

### The P-T286 reaction prefers a *cis*-holoenzyme mechanism

In order to test for *cis*-versus *trans*-holoenzyme autophosphorylation, we conducted biochemical reactions with CaMKII WT holoenzymes and GFP-tagged CaMKII K42M mutant holoenzymes (Figure 2A). The K42M mutation makes CaMKII kinase dead, as it prevents ATP binding^13,24,25^; the GFP-tag allows separation of the K42M mutant from CaMKII WT on a Western blot^26^ (Figure 2A). When CaMKII WT and GFP-CaMKII K42M are expressed separately, any detection of P-T286 into the kinase dead GFP-CaMKII K42M mutant (that cannot phosphorylate itself) must have occurred either via a *trans*-holoenzyme reaction (Figure 2A) or by subunit exchange followed by a subsequent *cis*-holoenzyme reaction. In our experimental setup, the separately expressed CaMKII forms are mixed immediately before the reaction, favoring detection of the *trans*-holoenzyme reaction on the kinase dead GFP-CaMKII K42M mutant (based on the time scale of subunit exchange and suppression of the exchange-promoting autophosphorylation at the secondary T305/306 sites under our reaction conditions^17,18,26^). Indeed, P-T286 was detected on the GFP-CaMKII K42M mutant, but only in reactions where it was mixed with CaMKII WT (Figure 2B,C). Thus, consistent with previous studies^6,19^, the P-T286 reaction can indeed occur by a *trans*-holoenzyme reaction. However, P-T286 on CaMKII WT was more readily detected also at lower holoenzyme concentrations (Figure 2B) and occurred on a much faster time scale in CaMKII WT (Figure 2C), indicating that the *cis*-holoenzyme reaction is much more efficient.

**Figure 2:**
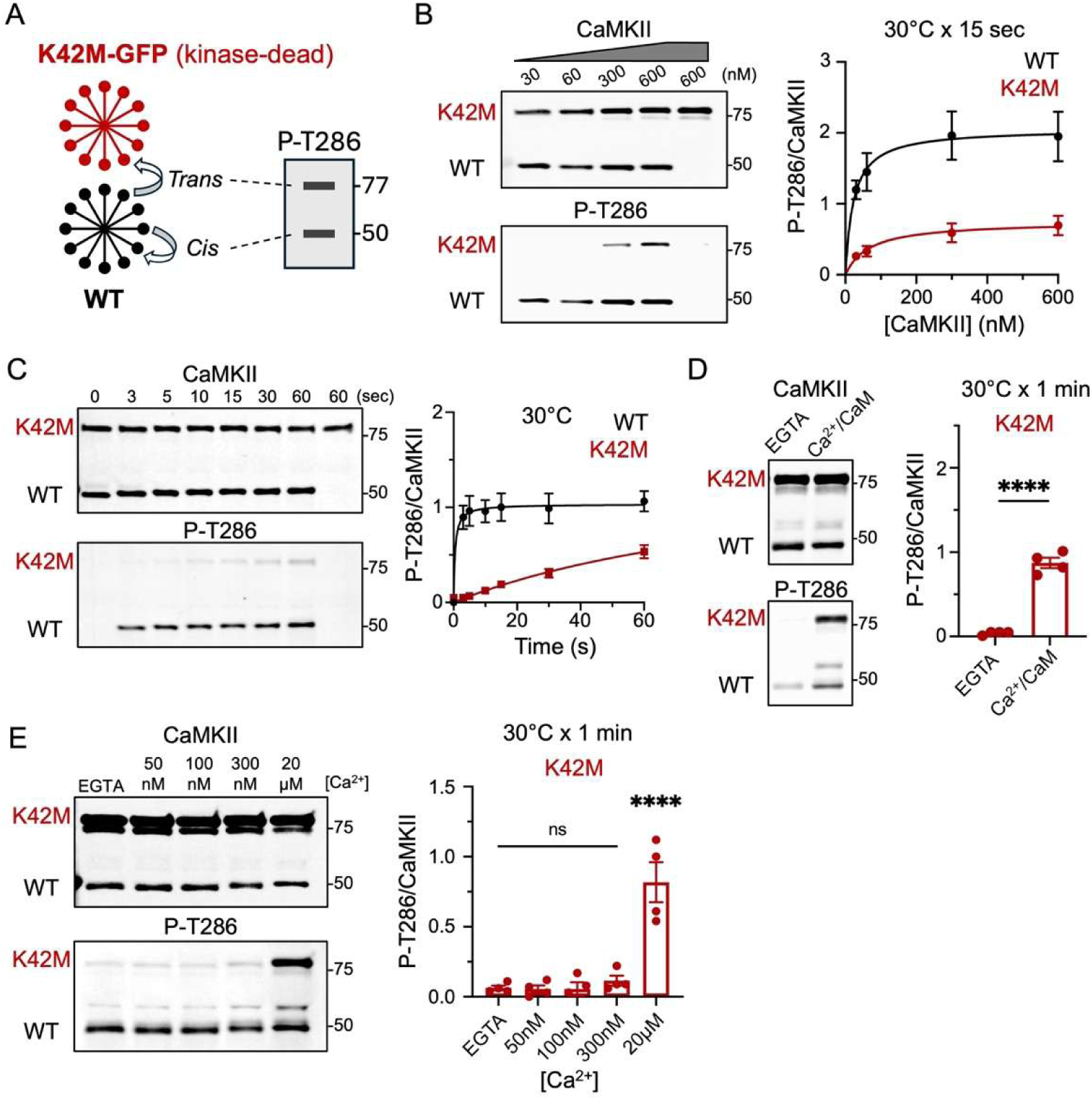
Trans-holoenzyme P-T286 does not occur at basal Ca2+ levels. Error bars indicate SEM in all panels. ****P<0.0001; ns, not significant. (A) Schematic of the assay: WT CaMKII (~50 kDa) mixed with kinase-dead GFP-tagged K42M (~77 kDa). Western blot detection of P-T286 on WT indicates *cis*-holoenzyme phosphorylation; on K42M indicates *trans*. Note that both the *cis*- and *trans*-holoenzyme P-T286 reaction has to occur between individual subunits. (B) Representative blots showing *in vitro* kinase reactions (30, 60, 300, and 600 nM kinase, 50 mM HEPES pH7.2, 1.5 mM Ca^2+^, 10 mM Mg^2+^, 1 mM ATP, 1 µM CaM) in which untagged-WT and mEGFP-K42M CaMKII constructs, isolated from HEK293T cells, were mixed in the reactions. Final lane represents the K42M-only negative control. Quantification of P-T286/CaMKII (N=4) shows that *trans*-holoenzyme P-T286 is more kinase concentration dependent. (C) Representative blots showing *in vitro* kinase reactions (10 nM kinase, 50 mM HEPES pH7.2, 1.5 mM Ca^2+^, 10 mM Mg^2+^, 1 mM ATP, 1 µM CaM) in which untagged-WT and mEGFP-K42M CaMKII constructs, isolated from HEK293T cells, were mixed in the reactions. Quantification of P-T286/CaMKII (N=4) shows that *cis*-holoenzyme P-T286 is more efficient than *trans*-holoenzyme P-T286. (D) Representative blots showing *in vitro* kinase reactions (30 nM WT and 300 nM K42M, 50 mM HEPES pH7.2, 1.5 mM Ca^2+^, 10 mM Mg^2+^, 1 mM ATP, 1 µM CaM) in which untagged-WT and mEGFP-K42M CaMKII constructs, isolated from HEK293T cells, were mixed in the reactions. WT was pre-phosphorylated for 1 min prior to mixing with K42M. Quantification of P-T286/CaMKII (N=4) indicates that *trans*-holoenzyme P-T286 cannot occur without Ca^2+^/CaM stimulation. (E) Representative blots showing *in vitro* kinase reactions (10 nM WT and 100 nM K42M, 50 mM HEPES pH7.2, 0 to 20 µM Ca^2+^, 10 mM Mg^2+^, 1 mM ATP, 1 µM CaM) in which untagged-WT and mEGFP-K42M CaMKII constructs, isolated from HEK293T cells, were mixed in the reactions. WT was pre-phosphorylated for 1 min prior to mixing with K42M. Quantification of P-T286/CaMKII (N=4) indicates that *trans*-holoenzyme P-T286 cannot occur at basal Ca^2+^ levels, even with increasing K42M concentration.

### The minor *trans*-holoenzyme P-T286 reaction requires Ca^2+^/Calmodulin

The initial reactions that demonstrated that P-T286 can occur as a *trans*-holoenzyme reaction were done in presence of 1.5 mM Ca^2+^ and 1 µM CaM (Figure 2B,C). Next, we decided to test if the *trans*-holoenzyme reaction is possible without Ca^2+^. While the *cis*-holoenzyme reaction cannot occur without Ca^2+^ (as demonstrated previously^6,7^), this does not necessarily apply to the *trans*-holoenzyme reaction. In order to test for *trans*-holoenzyme reaction without Ca^2+^, we first autophosphorylated CaMKII WT at P-T286 before mixing it with GFP-CaMKII K42M. This pre-reaction makes CaMKII WT “autonomous” (*i.e.*, active independently of Ca^2+^/CaM). However, P-T286 of the GFP-CaMKII K42M holoenzymes by the autonomous CaMKII WT holoenzymes was still detected only when the reaction contained Ca^2+^/CaM, but not when Ca^2+^ in the reaction was chelated with EGTA (Figures 2D and S3). Thus, similar as previously described for the *cis*-holoenzyme P-T286 reaction^6,7^, the *trans*-holoenzyme reaction requires Ca^2+^/CaM binding not only to activate the CaMKII subunit that acts as the kinase, but also to expose T286 for phosphorylation on the CaMKII subunit that acts as the substrate. While P-T286 can substitute for Ca^2+^/CaM in activating the subunit acting as the kinase, Ca^2+^/CaM still remains required for exposing T286 on the subunit acting as substrate. Thus, P-T286 cannot self-propagate in the absence of Ca^2+^, neither by the less favored *trans*-holoenzyme reaction nor by a *cis*-holoenzyme reaction after subunit exchange.

### The minor *trans*-holoenzyme P-T286 reaction does not occur at basal Ca^2+^ (≤ 100 nM)

Based on previous findings for the *cis*-holoenzyme P-T286 reaction^6,7^, our finding that the *trans*-holoenzyme P-T286 reaction also requires Ca^2+^ did not come as a surprise. However, this still leaves open questions about how much Ca^2+^ would be required, specifically if basal cellular resting Ca^2+^ levels would be sufficient. This is because sufficiency of resting Ca^2+^ levels would make additional Ca^2+^ stimulation unnecessary, which could enable de-facto self-propagation of P-T286 for all practical cell signaling purposes. Basal cellular resting Ca^2+^ levels are generally thought to be around 50 nM, with 100 nM being on the high end^27–30^. In addition to 50 nM and 100 nM Ca^2+^, we tested potential trans-holoenzyme P-T286 reactions also at higher Ca^2+^ concentrations: 300 nM (corresponding to the weak Ca^2+^ signals estimated after LTD stimuli) and 20 µM (corresponding to strong Ca^2+^ signals after LTP stimuli)^26,31^. Any significant *trans*-holoenzyme P-T286 in the GFP-CaMKII K42M mutant by autonomous CaMKII WT was detected only at LTP level Ca^2+^ concentration (20 µM), but neither at basal Ca^2+^ levels (50 or 100 nM) nor LTD Ca^2+^ levels (300 nM) (Figure 2E). Notably, even for P-T286 detection at 20 µM Ca^2+^, we had to increase the ratio of the GFP-CaMKII K42M substrate holoenzymes over the CaMKII WT kinase holoenzymes (10:1 in Figure 2E) as no P-T286 was detected when the 1:1 ratio from the previous panels was used (Figure S4). Thus, P-T286 cannot self-propagate by a *trans*-holoenzyme reaction, neither in the absence of Ca^2+^ nor at the basal cellular resting Ca^2+^ levels of 50-100 nM.

### The major *cis*-holoenzyme P-T286 reaction does not occur at basal Ca^2+^ either

While the *trans*-holoenzyme P-T286 reaction did not occur at basal cellular Ca^2+^ (Figure 2E), this did not rule the possibility for the more efficient *cis*-holoenzyme to occur under these conditions. In order to test this, we co-expressed the kinase dead GFP-CaMKII K42M mutant with the non-tagged autonomously active CaMKII T286D mutant. The co-expression enables integration into mixed holoenzymes^32,33^ and using the autonomous CaMKII T286D mutant circumvents the requirement of prior P-T286 of the subunits acting as kinase (Figure 3A), which was possible only in the mixing experiment that tested the trans-holoenzyme reaction, but not in the co-expression experiment to test the *cis*-holoenzyme reaction here. In this case, *cis*-holoenzyme P-T286 was detected not only in reactions with the Ca^2+^ levels mimicking LTP conditions (20 µM Ca^2+^) but also with lower Ca^2+^ levels mimicking LTD conditions (300 nM Ca^2+^) (Figure 3B). However, at basal Ca^2+^ levels (50 or 100 nM), no P-T286 was detected above background (Figure 3B). Thus, at the basal cellular resting Ca^2+^ levels, P-T286 does not self-propagate at all, neither by a *trans*-holoenzyme reaction nor by the more efficient *cis*-holoenzyme reaction (that was proposed to mediate such self-propagation after subunit exchange).

**Figure 3:**
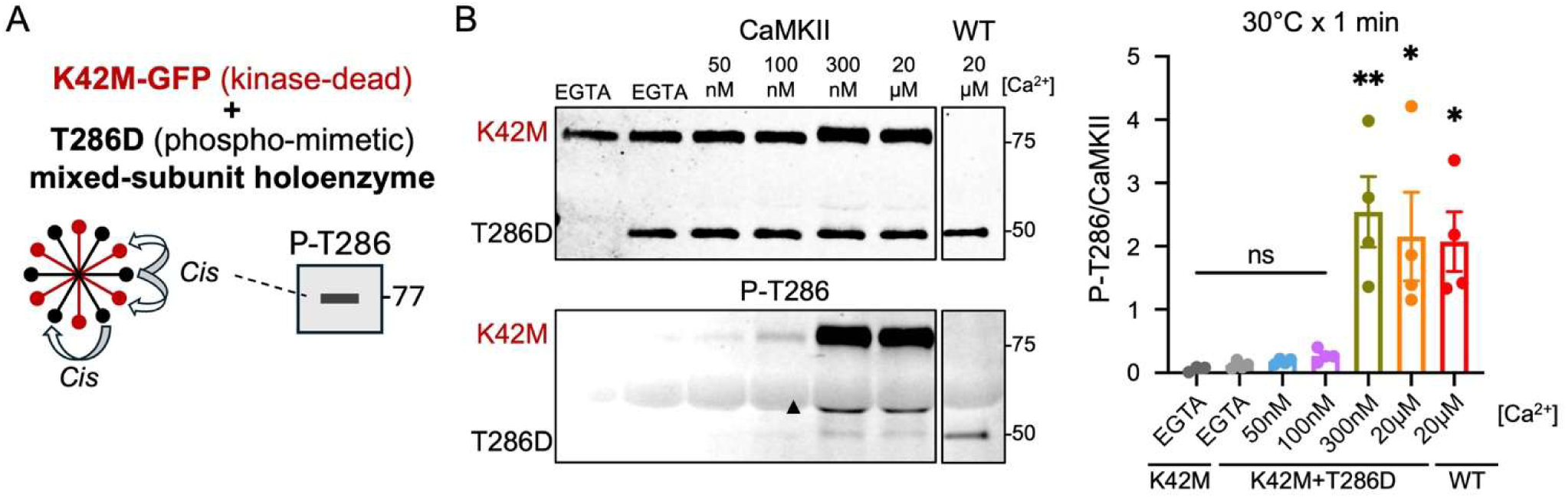
Cis-holoenzyme P-T286 does not occur at basal Ca2+ levels either. Error bars indicate SEM in all panels. *P<0.1; **P<0.01; ns, not significant. (A) Schematic of the assay: kinase-dead GFP-tagged K42M (~77 kDa) was co-expressed with phospho-mimetic T286D CaMKII (~50 kDa) to form mixed-subunit holoenzymes. Western blot detection of P-T286 on K42M indicates *cis*-holoenzyme phosphorylation between subunits driven by T286D subunits, as the T286D subunits can act only as kinase but not as substrate and the GFP-K42M subunits can act only substrate but not as kinase in the P-T286 reaction. (B) Representative blots showing *in vitro* kinase reactions (5 nM WT, 10 nM K42M, and 10 nM K42M+T286D, 50 mM HEPES pH7.2, 0 to 20 µM Ca^2+^, 10 mM Mg^2+^, 1 mM ATP, 1 µM CaM). K42M+T286D kinases were generated by co-expression of m-EGFP-K42M and untagged-T286D CaMKII constructs in HEK293T cells to produce mixed-subunit holoenzymes, and lysates were used for the reactions. WT and K42M were prepared separately as positive and negative controls. Triangle= non-specific bands. Quantification of P-T286/CaMKII (N=4) shows that *cis*-holoenzyme P-T286 cannot occur at basal Ca^2+^ levels.

### Maintenance of P-T286 is not required for maintenance of TBS-induced LTP

Our previous studies indicated that P-T286 is required during LTP induction only to enable efficient CaMKII binding to the NMDA-type glutamate receptor subunit GluN2B^13,14,34^ and that this requirement for P-T286 can be circumvented by ATP-competitive CaMKII inhibitors that directly enhance GluN2B binding^13,14^, at least for hippocampal LTP at the CA3 to CA1 synapse that is induced by high-frequency stimulation (HFS). Here, we tested the LTP at the same synapse, but after theta-burst stimulation (TBS) (Figure 4A), which is often regarded as more physiological. In slices from wildtype mice, addition of the ATP-competitive CaMKII inhibitor AS397 from 15 min before to 5 min after the TBS appeared to reduce the initial post-tetanic potentiation, but allowed development of normal levels of LTP (Figure 4B,C), consistent with requirement of CaMKII binding to GluN2B during LTP induction and requirement of enzymatic activity of the GluN2B-bound CaMKII not until the subsequent phase of LTP expression^13,14^. In contrast to wildtype, slices from P-T286-incompetent T286A mutant mice showed no significant TBS-induced LTP (Figure 4D,E). However, addition of AS397 from 15 min before to 5 min after TBS allowed development of LTP (Figure 4D,E), similar as seen before in the wildtype slices. Thus, after AS397 allowed the induction of LTP in the T286A slices, LTP expression and maintenance was normal despite the T286A mutation. Overall, our experiments show (i) that P-T286 cannot be maintained by self-propagation without additional Ca^2+^ stimuli and (ii) that such maintenance of P-T286 is not required for LTP maintenance.

**Figure 4:**
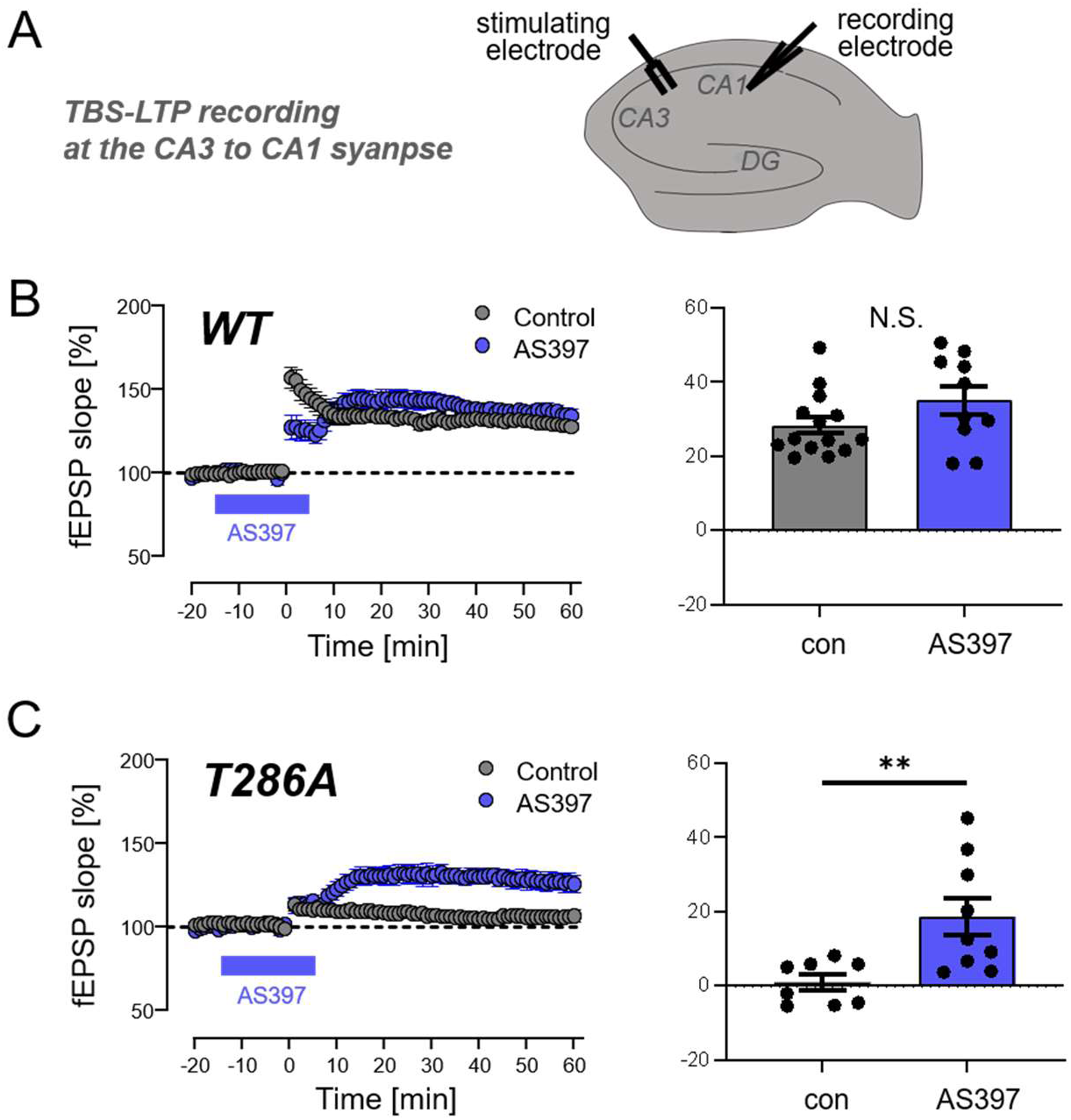
CaMKII P-T286 is required for LTP induction but not for LTP expression or maintenance. (A) Schematic of electrode placement in hippocampal slices for recoding of theta burst stimulation (TBS) induced LTP at the CA3 to CA1 synapse. (B) In wildtype slices, the ATP-competitive CaMKII inhibitor AS397 does not prevent LTP induction, with 10 uM AS397 applied from 15 min before to 5 min after TBS. While the immediate post-tetanic potentiation appears stunted, normal LTP develops after washout of the drug, consistent with LTP expression mediated by the autonomous activity of GluN2B-bound CaMKII. (C) While CaMKII T286A mutant mice are impaired for TBS-induced LTP, application of 10 uM AS397 from 15 min before to 5 min after TBS rescues LTP induction, with LTP expression after drug washout as in wild type mice. Then, after rescue of LTP induction by AS397, LTP is normally maintained in the T286A mutant mice, indicating that P-T286 is not required for LTP expression or maintenance.

## DISCUSSION

The results of this study show that the autonomously active CaMKII P-T286 state cannot self-propagate in absence of Ca^2+^ stimuli, neither by subunit exchange nor by *trans*-holoenzyme autophosphorylation. Although we corroborated evidence for both subunit exchange and for *trans*-holoenzyme autophosphorylation, neither mechanism can propagate or maintain P-T286 without additional Ca^2+^ stimuli. While previous studies had already predicted that Ca^2+^/CaM binding is required to make T286 accessible for phosphorylation and therefore that Ca^2+^ is required for the P-T286 reaction^6,7^, it has been argued that basal levels of cellular Ca^2+^ (i.e. 50-100 nm) could be sufficient, at least if the reaction is mediated by a neighboring P-T286 subunit that is autonomously active and does not require Ca^2+^/CaM binding for stimulation of activity^10^. However, our results indicate that even an autonomous P-T286 subunit can mediate T286 phosphorylation of another subunit only during Ca^2+^ stimulation and neither in the absence of Ca^2+^ nor at basal levels of Ca^2+^. While the level of Ca^2+^ stimulation required for a P-T286 reaction within a holoenzyme appeared to be lower than the level required for a reaction between holoenzymes, basal levels of Ca^2+^ were not sufficient for either.

Maintenance of P-T286 by self-perpetuation had long been proposed to mediate synaptic memory, particularly in context of LTP maintenance^5,10^. However, the results of this study not only argue against the proposed P-T286 self-perpetuation but also demonstrate that P-T286 is not required for LTP maintenance: When the requirement for P-T286 to promote GluN2B binding during LTP induction is circumvented by ATP-competitive CaMKII inhibitors that can directly enhance this GluN2B binding, LTP is normally maintained after inhibitor washout even in slices from T286A mutant mice. Similar results have been previously obtained with different ATP-competitive inhibitors in two different studies for LTP induced by high-frequency stimulation (HFS)^13,14^. Our results on LTP induced instead by theta-burst stimulation (TBS) support the proposed underlying mechanism even more clearly, as the P-T286 incompetent T286A mutant mice still showed post-tetanic potentiation after HFS^13,14^ but not after the TBS in this study. This lack of post-tetanic potentiation helps to better unmask the different early phases of LTP: Presence of the CaMKII inhibitor before and during the TBS resulted in the same lack of post-tetanic potentiation during the LTP induction phase that primes for subsequent LTP expression by allowing CaMKII binding to GluN2B. Then, washout of the CaMKII inhibitor allows enzymatic activity of the GluN2B bound CaMKII (which is made Ca^2+^-independent by this binding^35^) to phosphorylate nearby substrates to mediate LTP expression. For instance, CaMKII-mediated phosphorylation can disperse SynGAP from synapses^36,37^, which in turn allows synaptic accumulation of Tarp-γ8 and AMPA-type glutamate receptors (AMPARs)^38^. Then, Tarps and AMPARs can be trapped at synapses by an additional round of local CaMKII-mediated phosphorylation^39,40^. Importantly, after successful LTP induction and expression, LTP is maintained normally in the T286A mice, demonstrating that P-T286 is not required for the maintenance of LTP, neither when induced by HFS nor by TBS.

As P-T286 cannot self-perpetuate without Ca^2+^ stimulation, what then is the functional role of the other parts of the proposed underlying mechanism, *i.e.*, subunit exchange between holoenzymes and *trans*-holoenzyme autophosphorylation? While this current study corroborated both mechanisms, their functional consequences remain to be explored. For *trans*-holoenzyme autophosphorylation, we here corroborate both its existence^19^ and the fact that the *cis*-holoenzyme autophosphorylation is much faster^6,19,41^. An argument for the signaling relevance of *trans*-holoenzyme autophosphorylation is the extreme abundance of CaMKII. In contrast to the *cis*-holoenzyme reaction, the *trans*-holoenzyme autophosphorylation reaction is dependent on the CaMKII concentration^6,19,41^; however, at the high CaMKII concentrations within neurons, such a *trans*-holoenzyme reaction would be entirely feasible. Nonetheless, a remaining argument against a significant contribution to signaling is the reaction speed: With the extremely fast *cis*-holoenzyme reaction speed^41^, it is unclear what additional signaling role a subsequent much slower *trans*-holoenzyme reaction could fulfill. In other words, if any given Ca^2+^ stimulus is not sufficient to already trigger the fast *cis*-holoenzyme reaction, how could it be sufficient to instead mediate the slower *trans*-holoenzyme reaction? Of course, the current lack of a good answer to this question does not preclude the discovery of such an answer in the future.

For subunit exchange, an obvious cellular function that is directly suggested by this study is the maintenance of holoenzyme integrity following subunit damage. Here, we find that proteolytic degradation of CaMKII holoenzymes promotes redistribution of hub assemblies from predominantly 12-mers into alternative oligomeric states, including 14-mers and partially opened 12-mers. Open-ring assemblies were not observed in our previous structural analyses of intact CaMKIIα or CaMKIIβ holoenzymes^22,23^; however, similar intermediates have been observed in an engineered variant of CaMKIIβ holoenzymes and proposed to participate in hub assembly/disassembly pathways associated with subunit exchange^42^. The appearance of open-ring structures following trypsin treatment is consistent with the idea that removal of the kinase and/or linker domains destabilizes hub interfaces and enables access to intermediate states involved in hub remodeling. In this context, such structural plasticity of the CaMKII holoenzyme could provide a mechanism for selective removal and replacement of damaged subunits without requiring complete disassembly and re-synthesis of the holoenzyme. Such a maintenance function could be particularly advantageous for large signaling assemblies such as CaMKII, where localized damage to individual subunits might otherwise compromise the integrity of the entire complex. Importantly, the same structural plasticity that enables repair and renewal of damaged assemblies may also provide the mechanistic foundation for other forms of regulating cellular signaling that are yet to be discovered. Indeed, subunit exchange is also described to be promoted by acute signaling events that lead to autophosphorylation of other sites within the regulatory domain, T305/306^17,18^.

Although CaMKII P-T286 has been most famous for its long-proposed role in storage of information^8,10^, there also have been arguments for the alternative role in computation of information that is now further supported by this study (as we have reviewed previously^3,5,15^). Indeed, the P-T286 mechanism within the CaMKII holoenzyme is ideally suited for signal computation: The dual requirement of Ca^2+^/CaM in the P-T286 reaction^6,7^ enables the frequency detection by P-T286 within the CaMKII holoenzyme^33,43^, with additional regulatory crosstalk with GluN2B binding and additional autophosphorylation at T305/306^13,35,44–46^ enabling different types of Ca^2+^ stimuli to induce CaMKII-dependent LTP or LTD^2,26^ (as recently reviewed^5^). These computational functions and the results of the current study do not rule out additional functions of CaMKII also in information storage. However, such storage would have to be mediated by different mechanisms (such as the Ca^2+^-induced but persistent binding of CaMKII to GluN2B^35,47–49^) and not by the P-T286 that instead mediates the computation of synaptic Ca^2+^ signals.

## ACKNOWLEDGEMENTS

This work was supported by National Institutes of Health grants R01 AG067713 and R01 NS081248 (to K.U.B.), and R35 GM124779 and R01 EY030987 (to S.L.R.). We thank Dr. Claudia López and the staff of the OHSU Multiscale Microscopy Core for training and instrumentation access.

## AUTHOR CONTRIBUTIONS

F.Y.C., N.L.R., A.P.M., and S.J.C. performed experiments and analyses; F.Y.C., S.L.R., and K.U.B. conceived this study; K.U.B. wrote the initial draft and all authors contributed to the final manuscript.

## STAR METHODS

### Key Resources Table

| REAGENT or RESOURCE | SOURCE | IDENTIFIER |
| --- | --- | --- |
| **Antibodies** |  |  |
| Anti-CaMKIIα | Made in house | CBα2; RRID: AB_2533032 |
| Anti-CaMKIIα (against AA 448-460) | BD Biosciences | 611293 |
| Anti-CaMKII P-T286 | PhosphoSolutions | P1005-286; RRID: AB_2492051 |
| Goat anti-mouse AzureSpectra 700 | Azure Biosystems | AC2129 |
| Goat anti-rabbit AzureSpectra 800 | Azure Biosystems | AC2134 |
| **Chemicals, Peptides, and Recombinant Proteins** |  |  |
| Penicillin-Streptomycin | Gibco | 15070063 |
| DMEM media | Gibco | 11965-092 |
| cOmplete protease inhibitor cocktail (EDTA-free) | Roche | 1187380001 |
| Microcystin-LR | Calbiochem | 475815 |
| Fluorescent Blot Blocking Buffer | Azure Biosystems | AC2190 |
| Calmodulin | Made in house | N/A |
| Adenosine triphosphate (ATP) | Sigma | A2383 |
| Trypsin | Sigma | T4799 |
| Phenylmethanesulfonyl Sulfate | Sigma | P6729 |
| Uranyl Formate | SPI-Chem | 02545-AB |
| **Deposited Data** |  |  |
| Raw and analyzed data | This paper |  |
| **Experimental Models: Cell Lines** |  |  |
| HEK293T cells | ATCC | CRL-1573 |
| **Experimental Models: Organisms/Strains** |  |  |
| Mouse: wild type: C57BL/6 | Charles River Labs |  |
| Mouse: CaMK2A T286A |  |  |
| **Deposited data** |  |  |
| Raw and analyzed data | This paper | (Mendeley link to be added) |
| **Software and Algorithms** |  |  |
| Slidebook 2025 | Intelligent Imaging Innovations (3i) | RRID:SCR_014300 |
| Prism 11.0.1 | Graphpad | RRID: SCR_002798 |
| FIJI ImageJ | NIH | RRID:SCR_003070 |
| CryoSPARC | Structura Biotechnology Inc | V4.5.1 |

### LEAD CONTACT AND MATERIALS AVAILABILITY

Further information and requests for resources and reagents should be directed to and will be fulfilled by the Lead Contact, K. Ulrich Bayer. This study did not generate new unique reagents.

### EXPERIMENTAL MODEL AND SUBJECT DETAILS

All animal treatments were approved by the University of Colorado Institutional Animal Care and Use Committee, in accordance with NIH guidelines, and was done as described previously ^50^. Animals are housed at the Animal Resource Center at the University of Colorado Anschutz Medical Campus (Aurora, CO) and are regularly monitored with respect to general health, cage changes, and overcrowding.

### METHOD DETAILS

#### Material and DNA constructs

Material was obtained from Sigma, unless noted otherwise. Mutant CaMKII constructs were generated by PCR-based site-directed mutagenesis from WT CaMKII plasmids (mEGFP-tagged or untagged), which served as templates for amplification. All constructs were validated by Sanger sequencing.

#### In vitro phosphorylation assays

For *trans*-holoenzyme assays, WT and K42M CaMKII constructs were expressed in HEK293T cells separately. The lysates were prepared independently and mixed immediately prior to the reaction. For *cis*-holoenzyme assays, K42M CaMKII was co-expressed with T286D CaMKII in HEK293T cells, and lysates were used directly. Kinase reactions were performed in a reaction buffer containing 50 mM HEPES (pH7.2), 10 mM MgCl_2_, 1 mM ATP, and 1 µM CaM, with CaCl_2_ concentrations ranging from 0 to 20 µM or 1.5, mM as indicated in each figure. Total CAMKII concentrations (reported as kinase subunit concentration) ranged from 5 to 600 nM, depending on the experiment. Reactions were carried out at 30°C for the indicated time and were initiated by addition of reaction buffer to the lysate. Where indicated, WT CaMKII was pre-phosphorylated for 1 min prior to mixing with K42M. Reactions were terminated by adding western blot sample buffer containing 1.6 % SDS and 25 mM EDTA and heating at 95°C for 5 min.

#### Western analysis

Samples were boiled in Laemmli sample buffer for 5 min at 95°C prior to SDS-PAGE. Proteins were separated on 4-20% Criterion TGX precast polyacrylamide gels (Bio-Rad) and transferred to a fluorescence-optimized PVDF membrane at 24 V for 1.5 hours at 4°C. Membranes were blocked in Fluorescent Blot Blocking Buffer (Azure Biosystems) for 1 hour at room temperature, followed by incubation with primary antibodies overnight at 4°C, including anti-CaMKIIα (1:2000, CBα2; available at Invitrogen but made in house) and anti-P-T286 (1:3000; PhosphoSolutions). After washing, membranes were incubated with goat anti-mouse AzureSpectra 700 (1:6000; Azure Biosystems) and goat anti-rabbit AzureSpectra 800 (1:4000; Azure Biosystems) for 1 hour at room temperature. Blots were developed using an Azure 600 Western Blot Imager (Azure Biosystems) and analyzed by densitometry (FIJI ImageJ). Phospho-signal was normalized to total CaMKII levels.

#### Protein extracts

Whole homogenates of HEK cells were made in homogenization buffer including 50 mM PIPES pH 7.2, 10% glycerol, protease inhibitor cocktail and 1 mM DTT. The homogenates were cleared by centrifugation at 14,000 RPM at 4°C for 20 min. For all expression experiments, cells were harvested 48 h after transfection.

#### Trypsin Proteolysis Experiments

Purified, recombinant rat CaMKIIα (2.6 µM) was digested with 0.01 mg/ml Trypsin in a buffer containing 20 HEPES pH 7.4, 150 KCl, 1 EDTA as indicated. For analysis by EM, digestion was performed at room temperature, either overnight, then stopped with 5 mM PMSF or for 1 h, stopped with 5 mM PMSF then further incubated overnight. Kinase was then frozen at −80°C.

#### Negative-stain EM of full-length and trypsinized CaMKIIα

Full-length CaMKIIα, 1-hour trypsinized CaMKIIα, and overnight trypsinized CaMKIIα samples were prepared for negative-stain EM by dilution to ~100 nM subunit concentration in buffer containing 20 mM HEPES (pH 7.4), 150 mM KCl, and 1 mM EDTA. For each condition, 3 µL of sample was applied to freshly glow-discharged (15 mA, 1 min) 400-mesh copper EM grids (Ted Pella) coated with a continuous carbon support (~12 nm thickness). Excess sample was blotted with filter paper, grids were washed twice with ultrapure water, and stained with 0.75% (wt vol^-1^) uranyl formate (SPI-Chem), blotted with filter paper and dried with laminar airflow.

Negatively stained CaMKIIα samples were imaged on a 120 kV TEM (Tecnai iCorr, FEI) equipped with a CCD camera (AMT XR16). Micrographs were manually collected using AMT Image Capture Engine (v602.591j) at a nominal magnification of 30,000×, corresponding to a calibrated pixel size of 3.991 Å pixel^-1^, with a nominal defocus range of 1.7–2.2 µm. A total of 25 micrographs were collected for full-length CaMKIIα, 8 micrographs for 1-hour trypsinized CaMKIIα, and 10 micrographs for overnight trypsinized CaMKIIα.

Micrographs were imported into CryoSPARC (v4.5.1)^51^ for image processing and particle classification. Approximately 100 particles from each dataset were manually picked and subjected to reference-free 2D classification to generate templates for automated particle picking. Micrographs and particles were CTF corrected by phase flipping. Multiple rounds of 2D classification were performed to remove non-particle picks and poorly aligned particles. For the full-length CaMKIIα controls specimen, 1,396 particles were extracted using a box size of 76 pixels and subjected to focused 2D classification (K = 50) using a binary soft mask (153 Å diameter) centered on the hub domain region. For trypsinized CaMKIIα samples, 1,515 particles from the 1-hour digest and 2,206 particles from the overnight digest were classified using identical parameters except with a larger binary soft mask diameter of 165 Å to accommodate the expanded hub-only assemblies. The smaller mask applied to full-length CaMKIIα excluded peripheral kinase domains from contributing to alignment and classification.

All 2D classifications were performed using the following parameters: 12 Å maximum resolution, binary mask recentering (threshold = 0.1), five final full iterations, and ten online EM iterations. Resulting 2D class averages were manually grouped into morphological categories corresponding to sixfold symmetric hubs (12-mer), partially opened sixfold symmetric hubs (open 12-mer), sevenfold symmetric hubs (14-mer), or undefined classes that were excluded from analysis. The fraction of particles confidently assigned to each morphological category was calculated from the total number of classified particles.

#### Mouse hippocampal slice preparation

Hippocampal slices were prepared using 8-12 weeks old wildtype or CaMKII^ΔT286A^ mice. Isoflurane anesthetized mice were rapidly decapitated, and the brain was dissected in ice-cold high sucrose solution containing (in mM): 220 sucrose, 12 MgSO_4_, 10 glucose, 0.2 CaCl_2_, 0.5 KCl, 0.65 NaH_2_PO_4_, 13 NaHCO_3_, and 1.8 ascorbate. Transverse hippocampal slices (400 µm) were made using a tissue chopper (McIlwain) and transferred into 32°C artificial cerebral spinal fluid (ACSF) containing (in mM): 124 NaCl, 3.5 KCl, 1.3 NaH2 PO4, 26 NaHCO3, 10 glucose, 2 CaCl_2_, 1 MgSO_4_, and 1.8 ascorbate. Slices were allowed to recover for at least 1 h. All solutions were saturated with 95% O2/5% CO_2_.

#### Electrophysiology

All recordings and analysis were performed blind to genotype. A glass micro-pipette was used to record field excitatory post-synaptic potentials (fEPSPs) from the CA1 dendritic layer in response to stimulation in the Schaffer collaterals at the CA2 to CA1 interface using a tungsten bi-polar electrode. Slices were continually perfused with 30.5 ± 0.5°C ACSF at a rate of 3.5 ± 0.5 mL/min during recordings. Stimuli were delivered every 20 sec and 3 responses (1 min) were averaged for analysis. Data were analyzed using WIN LTP with slope calculated as the initial rise from 10 to 60% of response peak. Input/output (I/O) curves were generated by increasing the stimulus intensity at a constant interval until a maximum response or population spike was noted to determine stimulation that elicits 50% of maximum slope. Paired-pulse recordings (50 ms inter-pulse interval) were acquired from 50% max slope and no differences in presynaptic facilitation were seen in mutant slices. Stimulus intensity was set to 50% of the maximum response slope for TBS-induced LTP. A stable baseline was acquired for a minimum of 20 min prior to TBS (5 episodes of 0.1 Hz, each episode contains 10 stimulus trains of 5 pulses each at 100 Hz). For pharmacological treatment experiments, a stable baseline for a min of 10 min was acquired before treatment with AS397 for 15 min prior to TBS stimulation, to be washed out 5 min post TBS stimulation. The control condition is these experiments was just stimulation application and continuous recording. Change in slope was calculated as a ratio of the average slope of the 20 min baseline.

### QUANTIFICATION AND STATISTICAL ANALYSIS

All data are shown as mean ± SEM. Statistical significance is indicated in the figure legends. Statistics were performed using Prism (GraphPad) software. Imaging experiments were obtained and analyzed using SlideBook 2025 software. Western blots were analyzed using ImageJ (NIH). All data were tested for their ability to meet parametric conditions, as evaluated by a Shapiro-Wilk test for normal distribution and a Brown-Forsythe test (3 or more groups) or an F-test (2 groups) to determine equal variance. All comparisons between two groups met parametric criteria, and independent samples were analyzed using unpaired, two-tailed Student’s t-tests. Comparisons between three or more groups meeting parametric criteria were done by one-way ANOVA with specific post-hoc analysis indicated in figure legends. Comparisons between three or more groups with two independent variables were assessed by two-way ANOVA with Bonferroni post-hoc test to determine whether there is an interaction and/or main effect between the variables.

### DATA AND CODE AVAILABILITY

The datasets generated during this study are available through Mendeley.

