## Supplemental Figures for "CaMKII T286 autophosphorylation is not propagated at basal Ca^2+^ levels and is required only for the induction phase of LTP"

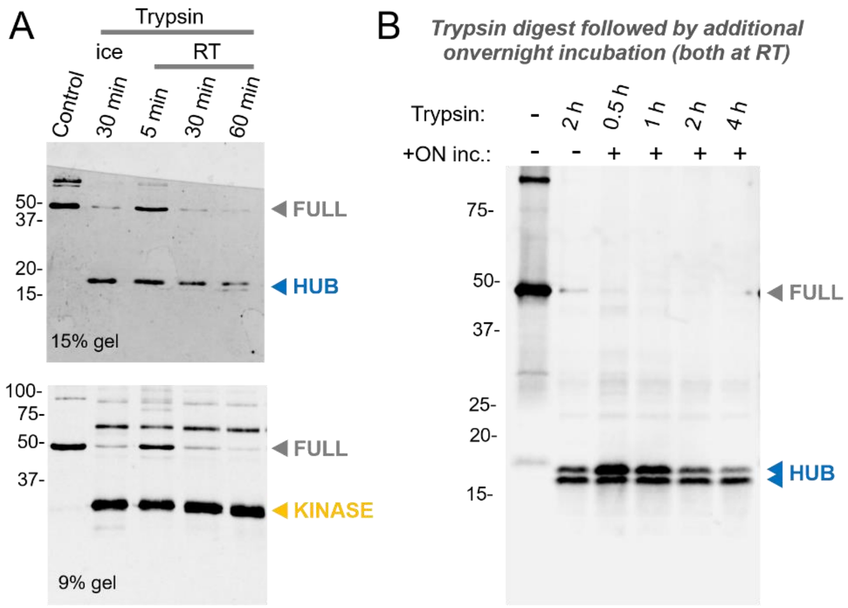


**Figure S1: Cleavage of CaMKII holoenzymes by trypsin to separate kinase and hub (association) domains.** Purified CaMKII was digested with trypsin under the conditions indicated. The digest was stopped with PMSF and the resulting fragments were analyzed by Western blot to verify digest prior to particle analysis by EM. **(A)** CaMKII holoenzymes are digested almost completely after 60 min digest with trypsin, Western blot detected fragments containing the association domain hub (upper blot; detected with the BD Biosciences antibody after PAGE on a 15% gel) or the kinase domain (lower blot; detected with the CBα2 antibody after PAGE on a 9% gel). **(B)** CaMKII holoenzyme digest at room temperature followed by continued overnight incubation at room temperature as indicated. The association domain is detected by the BD Biosciences antibody after PAGE on a 9% gel).

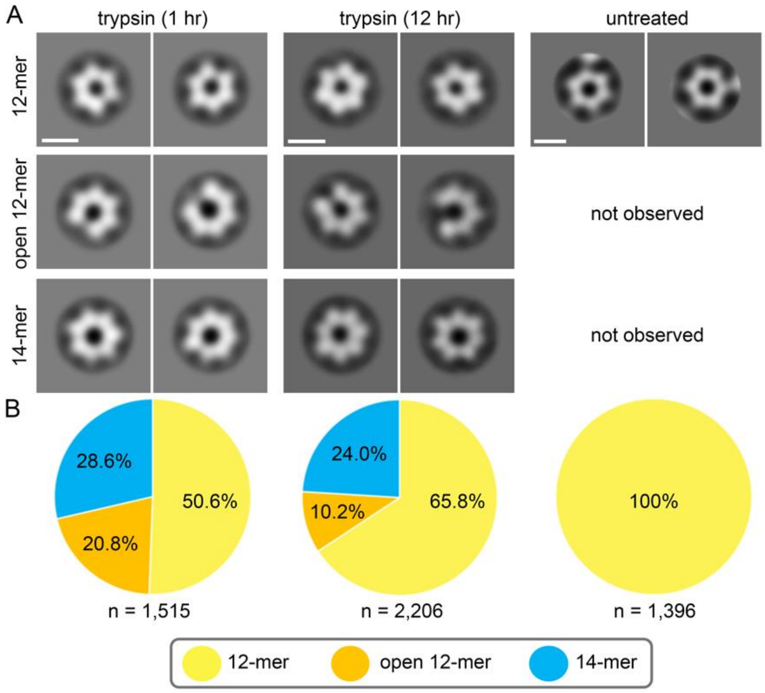


**Figure S2: EM analysis of CaMKII holoenzymes after extended trypsin digest** (12 h) shows qualitatively similar results as the 1 h digest followed by 11 h incubation shown in Fig. 1. Here, the results after 1 h versus 12 h are shown in direct comparison. **(A)** Representative focused 2D class averages obtained by negative-stain EM of hub-domain assemblies from untreated and trypsin-treated samples. Untreated particles classified exclusively as 12-mers, whereas trypsin-treated samples contained 12-mer, 14-mer, and open 12-mer assemblies. Scale bars, 10 nm. **(B)** Relative abundance of each hub-domain architecture determined by focused 2D classification. Numbers indicate the percentage of particles assigned to each class; *n* values are shown below each pie chart.

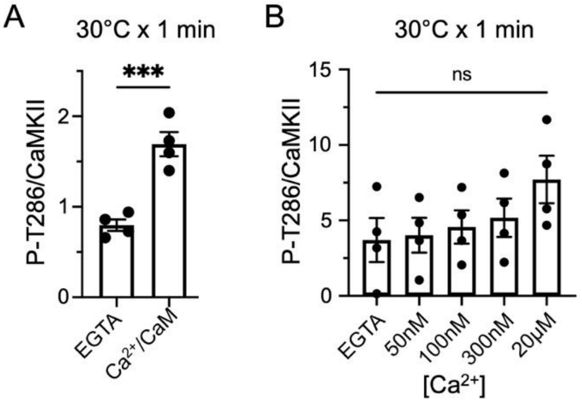


**Figure S3: WT CaMKII was pre-phosphorylated to induce autonomous activity** to test if the autonomous WT CaMKII could phosphorylate the kinase dead GFP-CaMKII K42M in absence of $Ca^{2+}$. The resulting P-T286 of GFP-CaMKII K42M is shown in Figure 2. This Figure shows the P-T286 of WT CaMKII, as quantified after Western analysis. Error bars indicate SEM in all panels. ***P<0.001; ns, not significant. **(A)** Quantification of P-T286/CaMKII (N=4) for WT CaMKII for the reactions shown in Figure 2D. Even after pre-phosphorylation, WT P-T286 levels were further increased during the $Ca^{2+}$/CaM-containing reaction. **(B)** Quantification of P-T286/CaMKII (N=4) for WT CaMKII for the reactions shown in Figure 2E. In this case, the mild apparent increase in P-T286 of the pre-phosphorylated WT CaMKII during the reactions containing some $Ca^{2+}$ was not statistically significant.

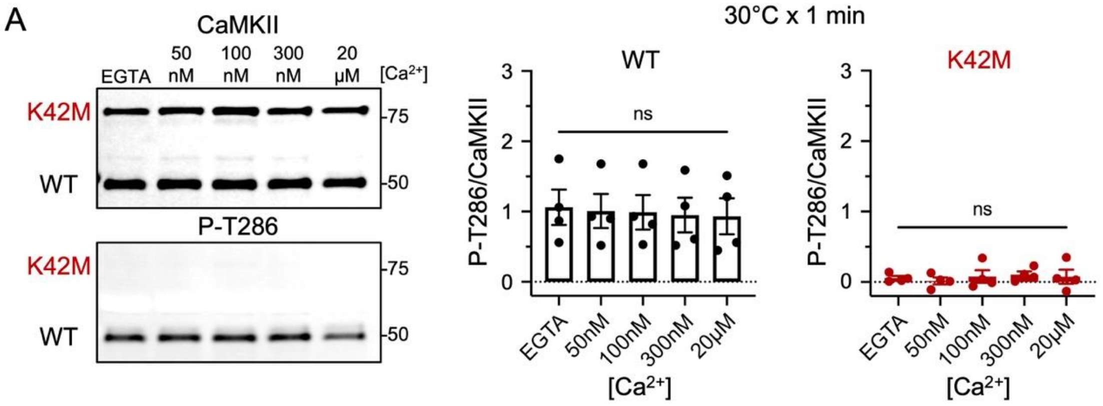


**Figure S4: *Trans*-holoenzyme P-T286 was undetectable at physiological $Ca^{2+}$ levels when the ratio of CaMKII K42M to WT was ~1:1.** Error bars indicate SEM in all panels. ns, not significant.
**(A)** Representative blots showing *in vitro* kinase reactions (10 nM WT and 10 nM K42M, 50 mM HEPES pH7.2, 0 to 20 μM $Ca^{2+}$, 10 mM $Mg^{2+}$, 1 mM ATP, 1 μM CaM) in which untagged-WT and mEGFP-K42M CaMKII constructs, isolated from HEK293T cells, were mixed in the reactions. WT was pre-phosphorylated for 1 min prior to mixing with K42M. **(B)** Quantification of P-T286/CaMKII (N=4) validates that *trans*-holoenzyme P-T286 was not detectable an any of the tested $Ca^{2+}$ concentrations when the ratio of CaMKII K42M to WT was low (~1:1).
